# LRRK2 Kinase Mediates Increased GCase Activity in Microglia in Response to Proinflammatory Stimuli

**DOI:** 10.1101/2025.10.10.681687

**Authors:** Emma J. MacDougall, Carol X.-Q. Chen, Eric Deneault, Zhipeng You, Narges Abdian, Thomas M. Durcan, Konstantin Senkevich, Ziv Gan-Or, Edward A. Fon

## Abstract

Variants in the *LRRK2* and *GBA1* genes are among the most common risk factors associated with Parkinson’s disease (PD). Both patients carrying PD-associated variants in *GBA1*, encoding lysosomal enzyme glucocerebrosidase (GCase), and a subset of non-carrier patients have been shown to have reduced GCase enzymatic activity, suggesting that reduced GCase activity may be a feature of both genetic and a subset of sporadic PD. However, the effect of PD-associated variants in *LRRK2*, encoding a serine/threonine kinase, on GCase activity remains controversial, with conflicting results in various tissues and cell types. Moreover, rare patients carrying both *GBA1* and *LRRK2* risk alleles seem to have a more benign disease course than carriers of *GBA1* variants alone, suggesting a complex interplay between these two genes in PD. Here we evaluate the effect of LRRK2 kinase activity on GCase activity in human induced pluripotent stem cell (iPSC)-derived microglia (iMGs), a PD-relevant brain cell type expressing high levels of LRRK2. Using CRISPR editing, isogenic control iPSC lines were generated to match PD patient-derived iPSC lines harbouring the *LRRK2* p.G2019S, p.M1646T, or p.N551K-p.R1398H protective haplotype variants. Whereas iMGs harbouring the p.M1646T variant, and the protective haplotype, respectively increased and decreased phosphorylation of canonical LRRK2 substrate, Rab10, GCase protein levels and activity were not altered in any of the *LRRK2* variant lines. Additionally, whereas pharmacological inhibition of LRRK2 kinase activity had no impact on GCase activity in iMGs under basal conditions, it attenuated the increase in GCase activity elicited in response to interferon γ (IFNγ) treatment. Moreover, GCase activity induced by IFNγ was reduced in PD risk LRRK2 p.M1646T iMGs and increased in p.N551K-p.R1398H protective haplotype iMGs compared to their isogenic corrected controls, congruent with their respective effects on LRRK2 kinase activity and PD risk. Thus, our data suggests a role for LRRK2 kinase activity in regulation of GCase activity in response to neuroinflammation.

## Introduction

Parkinson’s disease (PD) is marked by death of dopamine neurons in the substantia nigra pars compacta region of the midbrain, resulting in the characteristic motor symptoms of the disease.^1^ Current PD therapeutics are symptom-modifying, lose efficacy over time, and do not alter disease progression.^1^ Among the most common genetic causes and risk factors for PD are mutations in the *LRRK2* and *GBA1* genes.^2^

*LRRK2* encodes a dual-function GTPase – Ser/Thr kinase, leucine-rich repeat kinase 2 (LRRK2). *LRRK2* PD pathogenic variants, including the most common p.G2019S, lead to increased kinase activity *in vivo*.^3^ Additionally, LRRK2 harbours variants that alter PD risk. The p.M1646T variant increases risk, reaching signifiance in the most recent PD genome wide association study (GWAS), while the p.N551K-p.R1398H protective haplotype is associated with decreased risk of PD.^4–7^ The p.M1646T variant is associated with increased kinase activity and the protective haplotype with reduced kinase activity.^8–10^ Substrates of LRRK2 kinase activity include a subset of Rab GTPases.^11–13^ LRRK2 has been implicated in many different cellular processes; notably, along with downstream Rabs, in the autophagy lysosomal pathway (ALP).^14–20^ There is emerging evidence for a role of LRRK2 in immune function. LRRK2 is highly expressed in immune cells, and has been implicated in their response to inflammatory stimuli.^21–25^ Further, proper function of the ALP and response to lysosomal damage induced by pathogens are key to mediating the immune response.^14,26,27^ Additionally, a common non-coding genetic variant in the 5’ region of LRRK2, found to be associated with increased PD risk by GWAS, has been shown to increase LRRK2 expression in microglia but not other brain cell types.^28^

Biallelic variants in *GBA1*, the gene encoding lysosomal enzyme glucocerebrosidase (GCase), may lead to Gaucher disease (GD). In GD, a loss of GCase function results in varied and, at times, neurologic disease manifestations. Heterozygous *GBA1* variants have been associated to PD.^29–32^ Unlike GD, the mechanism of pathogenicity of *GBA1* variants in PD remains less clear; however reduced GCase activity has been reported in PD patients with and without *GBA1* variants.^33–36^ GCase activity has been shown to be increased in whole blood of carriers of the LRRK2 p.G2019S or p.M1646T variants, while the p.N551K-p.R1398H protective haplotype is nominally associated with decreased GCase activity.^7,33^ Conversely, decreased GCase activity has been observed in iPSC-derived dopamine neurons carrying LRRK2 hyperactive variants.^37^ Additional studies utilizing *LRRK2* knockout mice, *GBA1* mutant mice, or double *LRRK2* and *GBA1* mutant *Drosophila* also support an interaction between the two enzymes.^38–40^ However, assessment of GCase activity in PD peripheral blood mononuclear cells (PBMCs) showed no effect of LRRK2 kinase activity on GCase activity.^41^ Work from Kedariti and colleagues indicates that the outcome may be cell type dependent, observing a positive correlation between LRRK2 kinase activity and GCase activity in the majority of models surveyed.^42^ Moreover, the discordance in these reports could also stem from the use of varied methods to assess GCase activity.

Clinically, PD patients carrying *LRRK2* variants have, on average, a milder disease course than those carrying *GBA1* variants.^2^ Rare PD patients carrying both a *LRRK2* variant and *GBA1* variant have a less severe phenotype than those carrying a *GBA1* variant alone.^43–45^ This points to a potential protective effect of *LRRK2* variants in *GBA1*-PD, and bolsters the idea that LRRK2 activity and GCase activity are positively correlated. Both LRRK2 kinase activity and GCase activity are being pursued as therapeutic targets in PD;^2,46,47^ thus it is crucial to understand the interplay between these two enzymes, and which patients will benefit from therapies targeting one or the other.

Here we assess the impact of the LRRK2 p.G2019S, p.M1646T and p.N551K-p.R1398H variants on GCase activity in PD-patient derived human induced pluripotent stem cell (iPSC)-derived microglial cells (iMGs) – a PD relevant cell type expressing high levels of both LRRK2 and GCase. We confirm that the p.M1646T PD risk variant leads to increased LRRK2 kinase activity and the p.N551K-p.R1398H protective haplotype to decreased in kinase activity. However, we see no effect of any of these LRRK2 variants, or of pharmacological LRRK2 kinase inhibition, on GCase protein levels or lysosomal GCase activity under basal conditions. In contrast, treatment of iMGs with interferon γ (IFNγ) increased lysosomal GCase activity, in a manner that could be attenuated by LRRK2 kinase inhibition. Further, when treated with IFNγ, LRRK2 p.M1646T PD risk variant iMGs showed increased GCase activity, whereas the p.N551K-p.R1398H protective haplotype showed decreased GCase activity, when compared to their respective isogenic control iMGs. This suggests that LRRK2 kinase activity and GCase activity are positively correlated in iMGs under proinflammatory conditions.

## Results

### Generation of LRRK2 isogenic control iPSCs and differentiation to iMGs

iPSCs were generated by reprogramming PD patient derived PBMCs heterozygous for the LRRK2 p.G2019S, p.M1646T, and p.N551K-p.R1398H (protective haplotype) variants (Fig 1a). A previously generated healthy control (LWT) line was also used in this study.^48^ LRRK2 knockout (LKO) and isogenic control lines with correction of LRRK2 variants were generated using CRISPR/Cas9 editing. Two isogenic control lines were generated for the LRRK2 protective haplotype - one with correction of the p.N551K variant (p.N551K^CORR^-p.R1398H), and one with correction of the p.R1398H variant (p.N551K-p.R1398H^CORR^). Correcting each variant individually allows for dissection of the effects of each variant of the haplotype. iPSC lines were subjected to quality control measures including karyotyping, and assessment of expression of pluripotency markers (Fig. S1 & S2).

**Figure 1.**
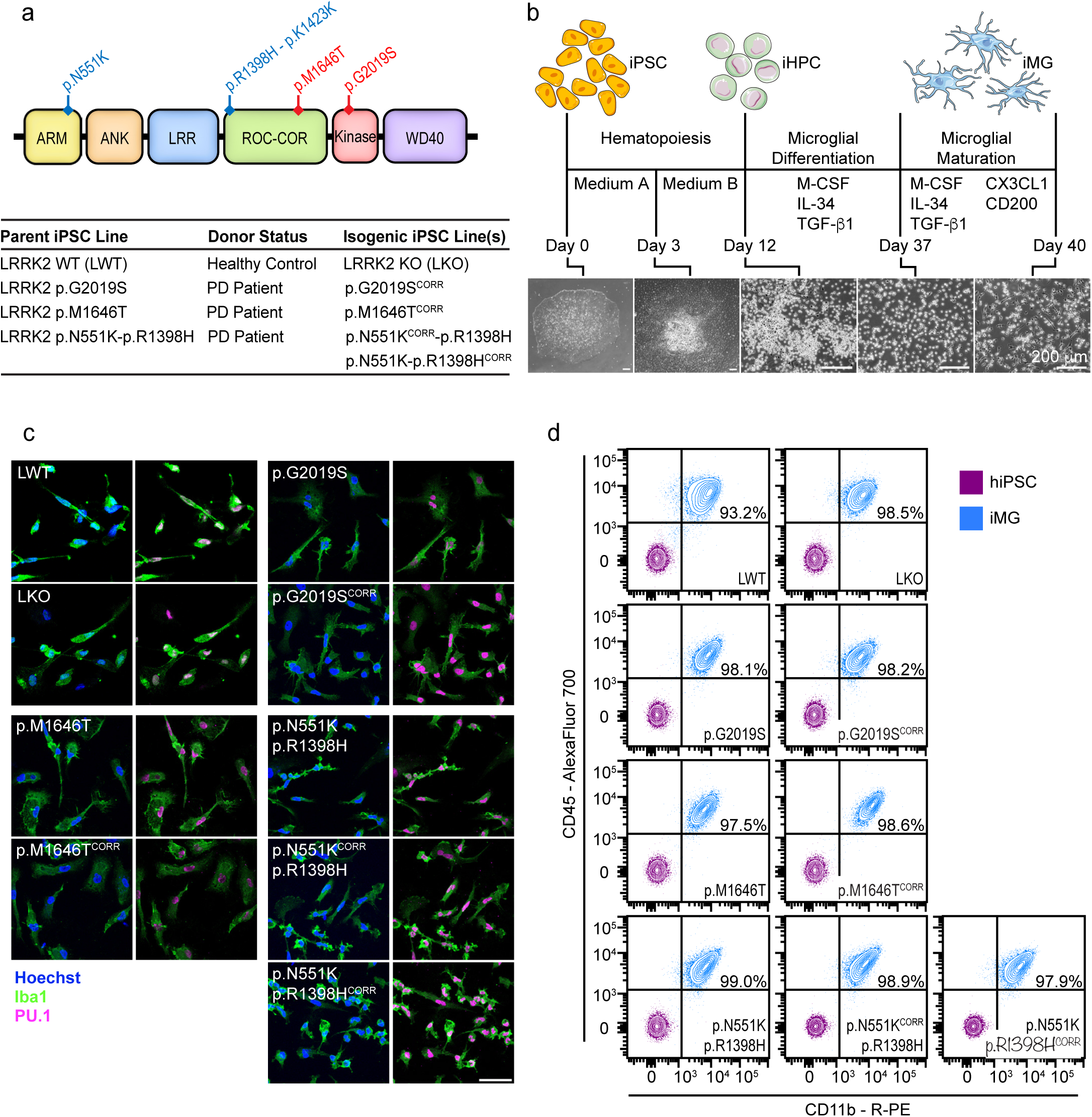
Differentiation and characterization of iMGs. **a** Schematic of LRRK2 domain structure and variants of interest. Variants predicted to increase kinase activity in red, and to decrease kinase activity in blue. Parental and matching isogenic iPSC lines used in this study. **b** Differentiation of iPSCs to iMGs. iPSCs are differentiated through mesoderm to produce hematopoietic progenitor cells (iHPCs) using Stem Cell Technologies STEMdiff™ Hematopoietic Kit (Day 0 – 12). iHPCs are further differentiated to iMGs via M-CSF, IL-34, and TGFβ1 supplementation (Day 12 – 37). Primitive iMGs are matured by the addition of CX3CL1 and CD200 (Day 37 – 40). Scale bars 200 μm. **c** iMGs express microglial markers Iba1 and PU.1, as detected by IF staining. Scale bar 50 μm. **d** iMGs from all lines are > 90% double positive for macrophage markers CD45 and CD11b by flow cytometry analysis.

All iPSC lines were differentiated into iMGs following the protocol published by McQuade and colleagues (Fig. 1b).^49^ Microglial identity was confirmed by assessing expression of canonical microglial (Iba1 and PU.1) and macrophage (CD11b and CD45) markers by immunostaining and flow cytometry, respectively (Fig 1c & d). iMGs generated from all lines were at least 90% double positive for CD11b and CD45 expression, indicating little to no contamination from non-myeloid lineage cells.

### LRRK2 variants alter kinase activity in iMGs

To assess the effects of the LRRK2 p.G2019S and p.M1646T risk variants, and the p.N551K-p. R1398H protective haplotype on LRRK2 kinase activity in iMGs, we performed western blotting using phospho-specific antibodies against LRRK2 and well-characterized LRRK2 substrates Rab10 and Rab12. Phosphorylation of LRRK2 was assessed both at its autophosphorylation site S1292, and its biomarker site S935. LRRK2 knockout iMGs were employed as a control for antibody specificity, and to establish levels of LRRK2-independent Rab phosphorylation. No significant differences in LRRK2 phosphorylation at either site were observed in LRRK2 variant iMGs compared to their isogenic controls (Fig S3a-c). A trend towards decreased LRRK2 S935 phosphorylation was observed with treatment using LRRK2 kinase inhibitor MLi-2, however was not significant except in LRRK2 p.M1646T iMGs (Fig S3c). MLi-2 treatment did however, lead to a significant decrease in pT73 Rab10 levels in all lines, excluding the knockout iMGs (Fig 2a & b). Despite the well-characterized increase in kinase activity associated with the LRRK2 p.G2019S variant, there was no difference in Rab10 phosphorylation in the LRRK2 p.G2019S iMGs compared to their isogenic controls – consistent with reports in mouse lung and kidney tissue and human neutrophils.^50,51^ In concordance with its risk variant status, and a previously published *in vitro* kinase assay,^8^ an increase in Rab10 phosphorylation was observed in the LRRK2 p.M1646T variant iMGs compared to isogenic control iMGs. As the p.R1398H variant is thought to drive the association of the protective haplotype to PD, and is known to impact both LRRK2 GTPase and kinase function,^52,53^ we chose to normalize both the protective haplotype iMGs and the p.N551K^CORR^ iMGs to the p.R1398H^CORR^ iMGs. Individual correction of both the p.N551K and p.R1398H variants of the LRRK2 protective haplotype increased phosphorylation of Rab10 (Fig 2a & b). Phosphorylation of Rab12 at S106 was also assessed. There was a trend towards decreased Rab12 phosphorylation with MLi-2 treatment, that reached significance in some lines, but no significant differences in LRRK2 variant iMGs compared to their isogenic controls (Fig S3a & d). Thus, the LRRK2 p.M1646T risk variant, and the p.N551K-p.R1398H protective haplotype respectively increase and decrease Rab10 phosphorylation in iMGs.

**Figure 2.**
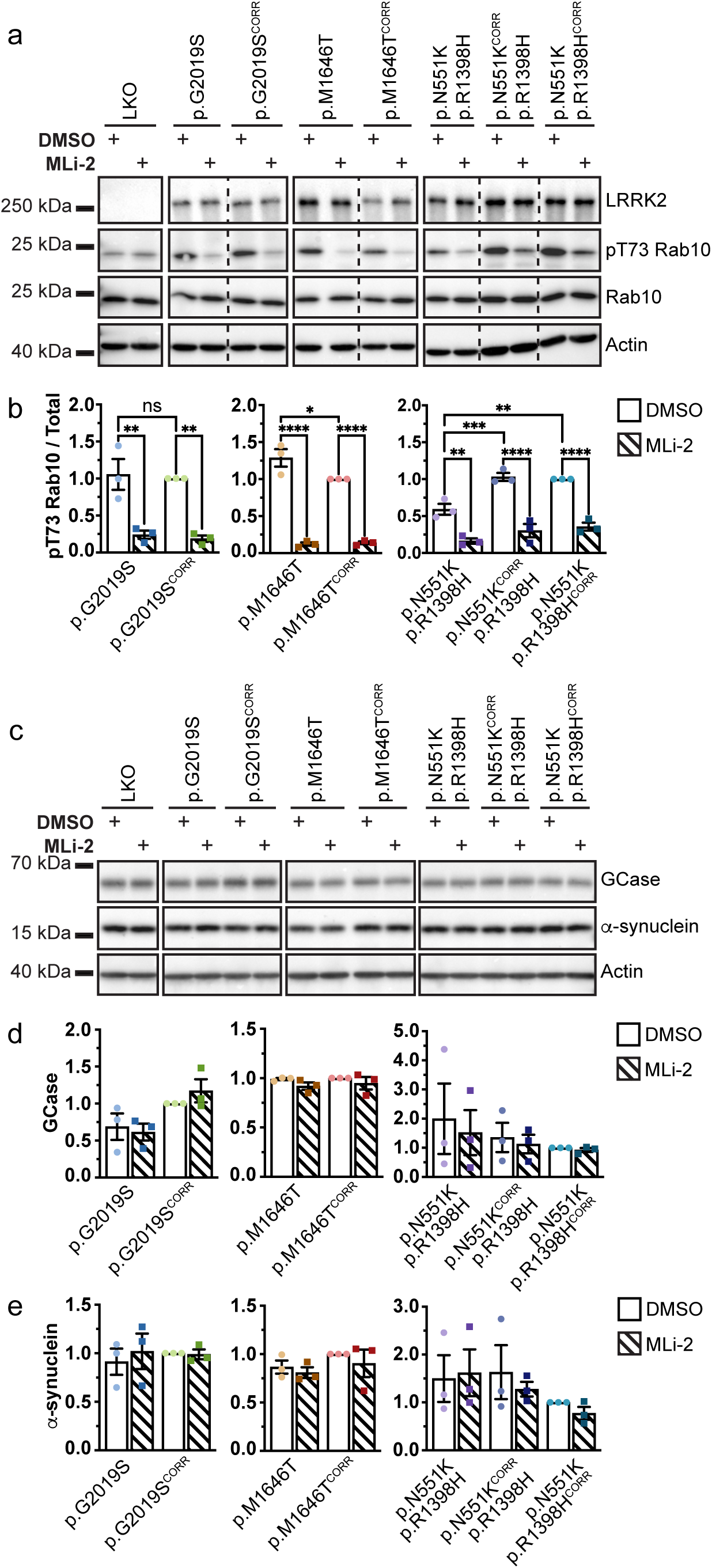
Rab10 phosphorylation is altered in LRRK2 variant iMGs, while GCase and α-synuclein levels are not **a** LRRK2 expression and Rab10 phosphorylation in LRRK2 variant iMGs with or without 6 hr 100 nM MLi-2 treatment as measured by WB. n = 3. **b** Quantification of WB of pT73 Rab10 normalized to total Rab10, levels normalized to isogenic control. **c** GCase and α-synucleinprotein levels in LRRK2 variant iMGs with or without 6 hr 100 nM MLi-2 treatment as measured by WB. n = 3. **d-e** Quantification of WB normalized to isogenic control. **d** Quantification of GCase. **e** Quantification of α-synuclein. One Way ANOVA with Bonferroni post-hoc test * p < 0.05, ** p < 0.01, **** p < 0.0001

### LRRK2 variants do not alter basal GCase or α-synuclein protein levels in iMGs

LRRK2 kinase activity has been reported to lead to altered levels of GCase protein,^42^ thus expression of GCase was assessed by western blotting. No significant differences in GCase protein levels were observed between LRRK2 variant iMGs and their isogenic controls. Additionally, LRRK2 inhibition had no effect on GCase levels (Fig 2c & d).

As LRRK2 variants and loss of GCase activity have both been shown to influence accumulation of α-synuclein,^54–56^ western blotting was used to assess changes in basal expression or accumulation of α-synuclein. There were no marked changes in α-synuclein protein levels in the LRRK2 variant iMGs or with LRRK2 kinase inhibition (Fig 2c & e). Overall, these data indicate that the changes in LRRK2 kinase activity or other potential effects associated with the p.G2019S, p.M1646T, and p.N551K-p.R1398H LRRK2 variants do not alter GCase or α-synuclein protein levels in iMGs under basal conditions.

### LRRK2 kinase activity has no effect on basal GCase activity in iMGs

To assess the effects of LRRK2 variants and their altered kinase activity on GCase activity in live cells, cleavage of the lysosome-specific fluorogenic GCase substrate 5-(Pentafluorobenzoylamino)Fluorescein Di-β-D-Glucopyranoside (PFB-FDGlu) was monitored by high-content microscopy (Fig 3a & Fig S4). The PFB-FDGlu fluorescence signal was not significantly different in LRRK2 variant or knockout iMGs compared to their isogenic controls at any point during the 140-minute incubation period (Fig 3b). Additionally, there were no differences in the rate of GCase activity, as represented by the slope of the linear portion of PFB-FDGlu fluorescence curves, in variant iMGs compared to their isogenic controls; or with use of two different LRRK2 inhibitors – MLi-2 or PF-475 (Fig 3c & Fig S5a). LRRK2 inhibition by PF-475 treatment in iMGs was confirmed by a reduction in Rab10 phosphorylation observed via western blot (Fig S5b). Mean total lysotracker area per cell was quantified as a proxy for lysosomal content, and despite implication of LRRK2 in lysosomal regulation, no differences were observed between LRRK2 variant iMGs and their isogenic controls (Fig 3d), or upon LRRK2 inhibition (Fig S5c). Taken together, these data indicate that there is no effect of LRRK2 kinase activity on GCase activity or gross lysosomal content in iMGs under basal conditions.

**Figure 3.**
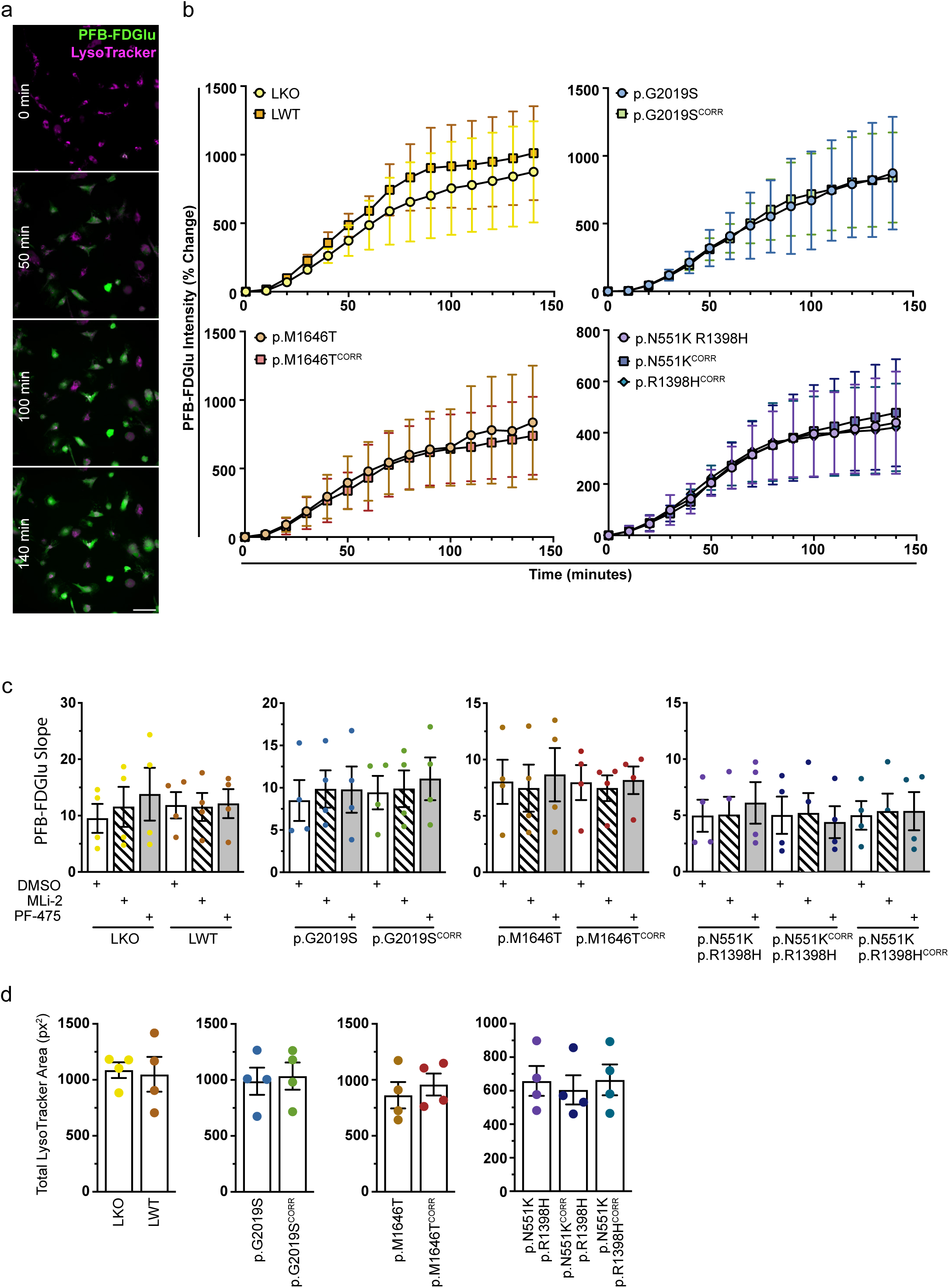
Lysosomal GCase activity is unaltered in LRRK2 variant iMGs under basal conditions **a** Representative images of LWT iMGs stained with lysotracker deep-red and PFB-FDGlu at 0, 50, 100 and 140 minutes after dye-loading. Scale bar 50 μm. **b** Percent change in PFB-FDGlu fluorescence. n = 4. **c** Slope of PFB-FDGlu fluorescence curves with or without 6 hr 100 nM MLi-2 or 500 nM PF-475 treatment. **d** Mean total lysotracker fluorescence area per cell (px^2^) at baseline (0 minutes). **c-d** Repeated Measures One Way ANOVA Tukey post-hoc test * p < 0.05, ** p < 0.01, **** p < 0.0001

### LRRK2 kinase inhibition reduces GCase activity in iMGs in response to proinflammatory stimuli

As neuroinflammation is a hallmark of PD, and LRRK2 expression levels are known to be increased in response to inflammatory stimuli,^22^ the impact of IFNγ treatment on GCase activity in iMGs was assessed. In healthy control (LWT) iMGs IFNγ treatment led to an increase in GCase activity, reflected by an increased slope of PFB-FDGlu fluorescence (Fig 4a & c & Fig S6). This is consistent with published data reporting increased GCase activity in PBMCs following IFNγ treatment.^41^ Interestingly, pretreatment of iMGs with MLi-2 followed by co-treatment with IFNγ and MLi-2 partially reversed the increase in GCase activity. The complete loss of PFB-FDGlu fluorescence upon treatment with lysosomal GCase inhibitor (CBE) demonstrates the specificity of the assay (Fig 4b-c). Additionally, no changes were observed in mean total lysotracker area of these cells (Fig 4d). Western blotting was performed to further assess lysosomal content and LRRK2 activity in iMGs upon IFNγ stimulation. In line with previous studies,^21,22^ LRRK2 expression, phosphorylation, and downstream Rab10 phosphorylation were increased in response to IFNγ treatment. Increased LRRK2 phosphorylation at S935 and Rab10 phosphorylation could both be blocked by MLi-2 treatment (Fig 5a-d). There were no marked changes in GCase or LAMP1 protein levels upon IFNγ or MLi-2 treatment (Fig 5a, e-f), indicating that the LRRK2-dependent increase in GCase activity observed under these conditions is unlikely to be driven by increased GCase protein or gross lysosomal content. Increases in lysosomal cathepsin activity have also been reported to occur in response to IFNγ treatment,^41^ thus global lysosomal proteolytic capacity was assessed by monitoring cleavage of fluorogenic protease substrate DQ Red BSA. No significant changes in global lysosomal proteolytic capacity were observed in response to IFNγ or MLi-2 treatment in LWT iMGs (Fig 5g & h). As a lysosomal enzyme, GCase functions optimally at a low pH (∼4.5-5.9)^57–59^. Both LRRK2 kinase hyperactivity and inflammatory stimuli have been reported to influence lysosomal pH.^17,60^ To assess if the IFNγ stimulated, LRRK2-dependent, increase in GCase activity that we observe is driven by a change in lysosomal pH we employed the pH-sensitive lysosomal-targeted pHLys Green probe. Increased fluorescence intensity of this probe is indicative of increased lysosomal acidity (lower pH). A subtle but significant increase in lysosomal acidity was observed after 24 hours of IFNγ treatment. However, LRRK2 inhibition via MLi-2 had no effect on this change in pH (Fig 5i & j). Together these data suggest that LRRK2 kinase signaling is involved in mediating an increase in GCase activity in response to inflammatory stimuli, independent of gross changes in lysosomal content, degradative capacity, and pH, or GCase protein levels.

**Figure 4.**
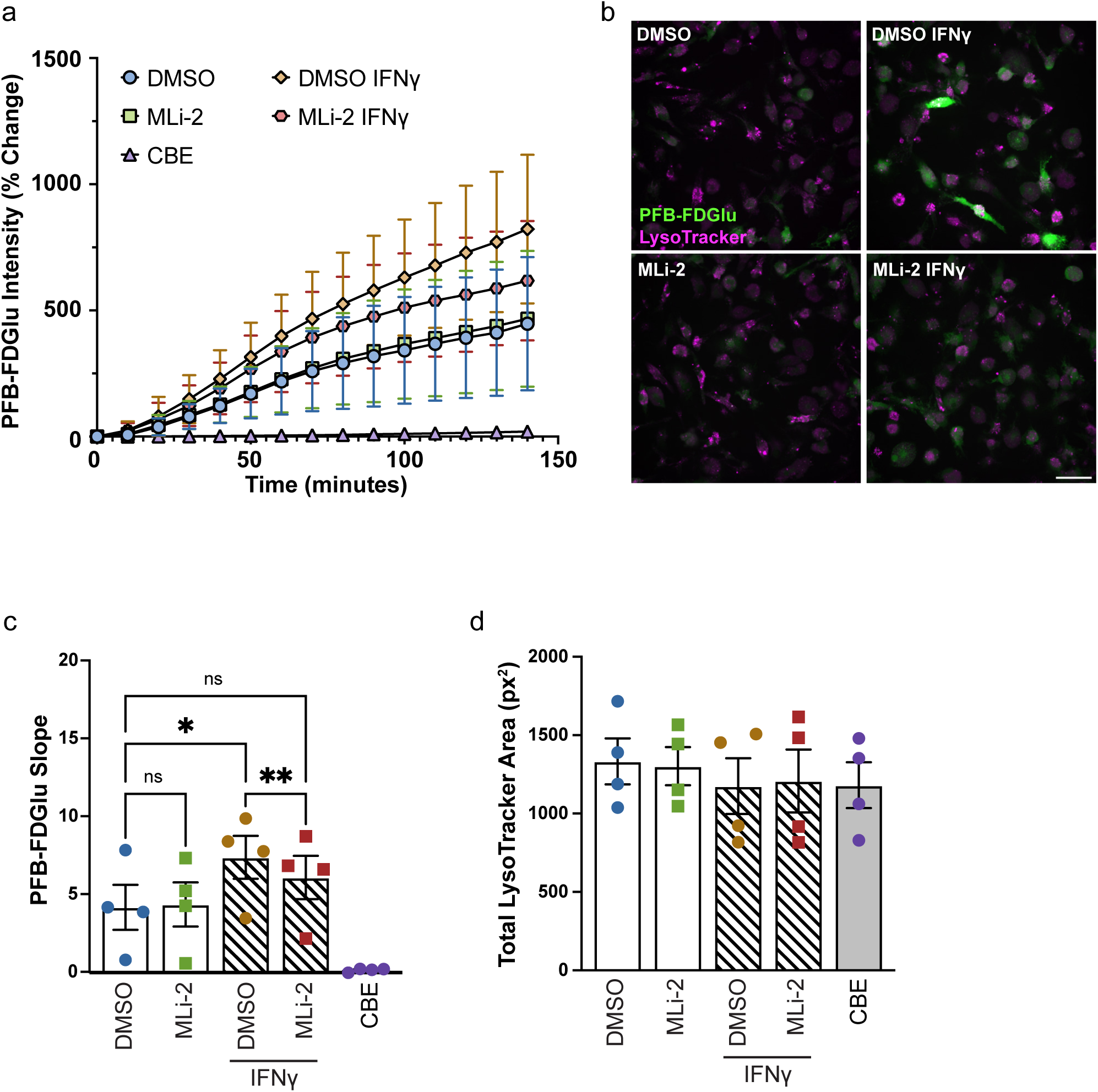
Lysosomal GCase activity is regulated by LRRK2 kinase activity under inflammatory stimulus **a** Percent change in PFB-FDGlu fluorescence in LWT iMGs with and without 24 hour 20 ng/mL IFNγ and 25 hour 100 nM MLi-2 treatment. n = 4. **b** Representative images of LWT iMGs, with and without IFNγ and MLi-2 treatment, stained with lysotracker deep-red after 140 minutes PFB-FDGlu incubation. Scale bar 50 μm. **c** Slope of PFB-FDGlu fluorescence. **d** Mean total lysotracker fluorescence area per cell (px^2^) at baseline (0 minutes). **c-d** Repeated Measures One Way ANOVA Tukey post-hoc test * p < 0.05, ** p < 0.01, **** p < 0.0001

**Figure 5.**
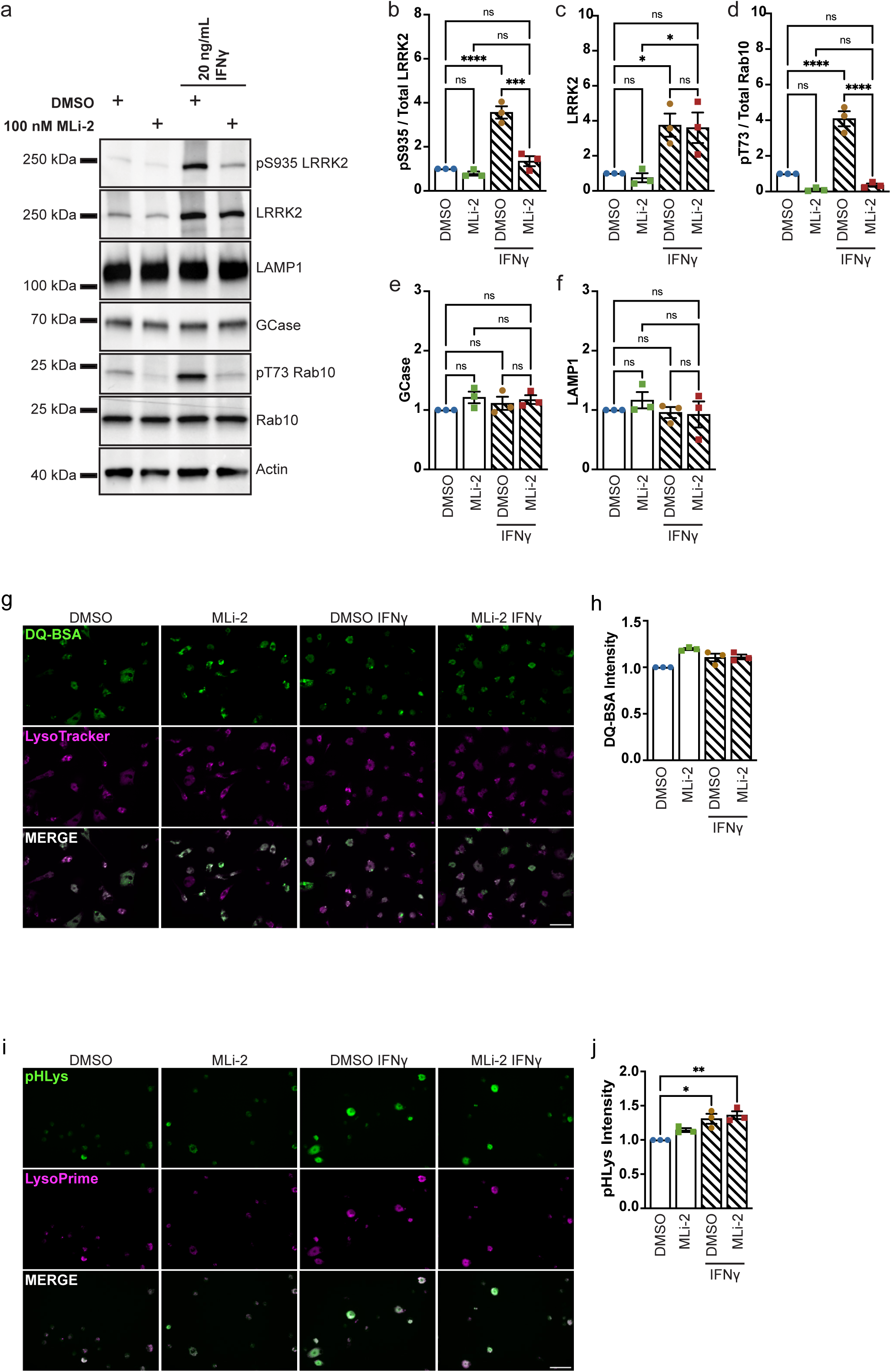
Inflammatory stimulus does not induce LRRK2-dependent changes in GCase protein expression, lysosomal proteolytic capacity, or pH. **a** Western blot analysis to assess LRRK2 levels and activity and lysosomal content in LWT iMGs with and without 24 hour 20 ng/mL IFNγ and 25 hour 100 nM MLi-2 treatment. n = 3. **b-f** Quantification of western blot. **b** Quantification of pS935 normalized to total LRRK2. **c** Quantification of LRRK2. **d** Quantification of pT73 normalized to total Rab10. **e** Quantification of GCase. **f** Quantification of LAMP1. **g** Representative images of DQ Red BSA signal in LWT iMGs following IFNγ and MLi-2 treatment 24 hours after dye-loading. **h** Mean DQ Red BSA fluorescence intensity normalized to mean total lysotracker area (px^2^) per cell, normalized to DMSO treated control, 24 hours after dye-loading. n = 3. **i** Representative images of pHLys Green signal in LWT iMGs following IFNγ and MLi-2 treatment. **j** Mean pHLys Green fluorescence intensity normalized to mean total LysoPrime Deep Red area (px^2^) per cell, normalized to DMSO treated control. n = 3. **b-i** One Way ANOVA with Bonferroni post-hoc test * p < 0.05, ** p < 0.01, **** p < 0.0001.

### PD-associated LRRK2 variants modulate GCase activity in iMGs in response to proinflammatory stimuli

We hypothesized that the LRRK2 kinase dependent increase in GCase activity observed in response to IFNγ treatment may reveal differences in GCase activity driven by PD-associated LRRK2 variants. Thus, we assessed GCase activity in LRRK2 variant iMGs and their isogenic controls after 24 hrs of IFNγ treatment, with or without pre- and co-treatment with MLi-2 (Fig 6). GCase activity was increased in all iMGs treated with IFNγ compared to non-treated controls, and consistent with our results in LWT cells, there was a trend for this increase to be attenuated by MLi-2 treatment – although only reaching significance in p.G2019S and p.M1646T variant iMGs (Fig 6c). In line with previous findings,^7^ IFNγ-treated p.M1646T iMGs exhibited increased GCase activity - illustrated by an increased slope of PFB-FDGlu fluorescence, compared to isogenic control iMGs. Additionally, correction of the p.N551K variant, but not the p.R1398H variant, of the LRRK2 protective haplotype led to an increase in GCase activity after IFNγ treatment. No effect of the LRRK2 variants or IFNγ treatment was observed on lysosomal content, represented by total lysotracker area (Fig 6d). This indicates that in p.M1646T and protective haplotype iMGs LRRK2 activity and GCase activity are positively correlated, with the change in GCase activity being driven by the p.N551K variant in the latter case. The LRRK2 p.G2019S variant has been reported to both increase and decrease GCase activity. Here we were unable to detect changes in GCase activity in these cells compared to their isogenic controls, even under IFNγ stimulation.

**Figure 6.**
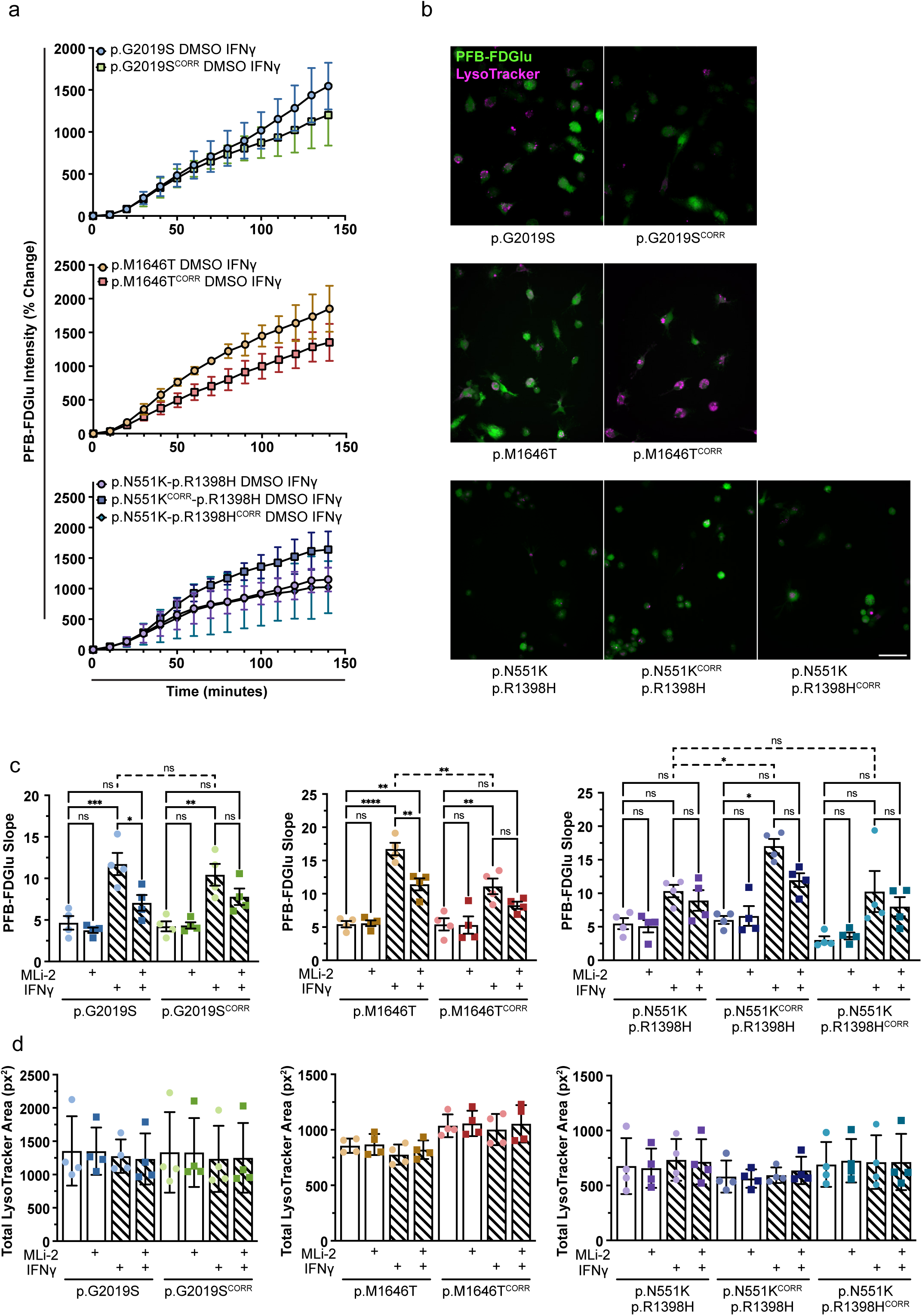
LRRK2 variants impact GCase activity under inflammatory stimulus. **a** Percent change in PFB-FDGlu fluorescence in LRRK2 variant and isogenic control iMGs upon 24 hour 20 ng/mL IFNγ and 25 hour DMSO treatment. n = 4. **b** Representative images of LRRK2 variant and isogenic control iMGs treated with IFNγ and DMSO 140 minutes after dye-loading. **c** Slope of PFB-FDGlu fluorescence in LRRK2 variant and isogenic control iMGs upon 24 hour 20 ng/mL IFNγ and 25 hour 100 nM MLi-2 treatment. **d** Mean total lysotracker fluorescence area per cell (px^2^) at baseline (0 minutes). **c-d** Repeated Measures One Way ANOVA Tukey post-hoc test * p < 0.05, ** p < 0.01, **** p < 0.0001

## Discussion

There is growing evidence for an interaction between LRRK2 and GCase activity. However, there is little agreement on the outcome of this interaction and how it may manifest in the context of PD. The aim of this work was to assess the effect of the LRRK2 p.G2019S, p.M1646T, and p.N551K-p.R1398H variants, all previously associated with altered GCase activity,^7,33,37^ on LRRK2 kinase activity and on GCase activity in iPSC-derived microglial cells. We demonstrate, for the first time in human PD-patient derived microglia, that in response to proinflammatory stimuli, LRRK2 kinase and GCase activities are positively correlated.

Phosphorylation of Rab10 was not increased in p.G2019S iMGs compared to isogenic control iMGs. This may be due to a subtle increase in kinase activity resulting from the p.G2019S mutation – altered Rab10 phosphorylation may be more readily detected using higher sensitivity methods such as mass spectrometry or mesoscale detection assays. These subtle changes may also be amplified by increased LRRK2 expression in response to inflammatory stimuli.^61^ Alternatively, there is evidence for differential phosphorylation of Rabs by various LRRK2 hyperactive variants across cell and tissue types.^50,51^ Consistent with a previously published *in vitro* kinase assay and its risk variant status,^8^ Rab10 phosphorylation was increased in p.M1646T iMGs. Like the p.R1441C/G/H and p.Y1699C PD pathogenic variants, the p.M1646T variant falls in the ROC-COR GTPase domain (Fig 1a), which is well-known to influence the kinase activity of LRRK2.^62^ We confirm, in iMGs, previous reports of decreased kinase activity associated with the LRRK2 protective haplotype.^10,52^ Interestingly, we find that correction of either the p.N551K or p.R1398H variant leads to increased phosphorylation of Rab10 – indicating that both variants can contribute to decreased kinase activity of the haplotype. The p.R1398H variant, localized in the GTPase domain, has been shown to be the primary driver of the association of the protective haplotype to PD, and to have increased GTPase activity and reduced kinase activity.^52,53^ An effect of the p.N551K variant alone on LRRK2 kinase activity has not been previously described. The p.N551K variant is localized to the N-terminal armadillo repeat (ARM) domain of LRRK2, which has recently been implicated in binding of LRRK2 to Rabs. Binding of Rab29 or Rab12 to the ARM domain drives membrane recruitment and activation of LRRK2,^63,64^ providing a potential mechanism for the altered kinase activity of the p.N551K variant *in vivo*.

Despite changes in kinase activity, the LRRK2 variants assessed had no effect on GCase protein levels or activity in iMGs under basal conditions. Likewise, LRRK2 KO and kinase inhibition had no effect on GCase expression or activity. LRRK2 kinase inhibition has been reported to have no effect on GCase activity, to increase GCase activity, and to decrease GCase activity across different studies and models.^37,39,41,42^ Notably, we employed 6 or 24 hour MLi-2 or PF-475 treatments (sufficient for dephosphorylation of Rab10 in iMGs. Fig 2a & b, Fig S3a, Fig 5a & d), whereas longer treatment periods (3-12 days) were utilized in studies where LRRK2 kinase inhibition led to increased GCase activity.^37,39^ Transcriptional changes induced by LRRK2 kinase inhibition over multiple days may underlie the altered GCase activity observed, and the same changes may be less likely to occur during the inhibition periods used here. Use of different methods of measuring GCase activity and cell models could also contribute to the inconsistencies in the effect of LRRK2 inhibition.

The lysosome is known to play a key role in mediating the immune response, but there is no consensus as to how the lysosome responds under IFNγ treatment.^65^ We find that GCase activity is increased with IFNγ stimulation, and that this increase can be blocked by LRRK2 kinase inhibition. Similarly, Wallings and colleagues report increases in GCase activity and pan-cathepsin activity in PBMCs stimulated with IFNγ; although these increases were not sensitive to LRRK2 kinase inhibition.^41^ Taken together these data could point to a global increase in lysosomal degradative capacity in response to IFNγ. However, using a pan-lysosomal protease substrate we did not observe a change in global lysosomal proteolytic activity in iMGs treated with IFNγ. IFNγ treatment did induce an increase in lysosomal acidity, but this was not attenuated by LRRK2 kinase inhibition. Moreover, we did not observe changes in gross lysosomal content in iMGs, as measured by total lysotracker area and LAMP1 protein levels, in response to IFNγ – thus it is unlikely that the LRRK2-driven increase in GCase activity is mediated by increased lysosomal mass, degradative capacity, or altered pH. GCase protein levels remained unchanged in response to IFNγ treatment, therefore the increased GCase activity observed is not driven by increased protein expression, whereas LRRK2 levels and Rab10 phosphorylation are increased upon IFNγ treatment. Whether increased GCase activity is driven solely by the increased expression of LRRK2, or by some other LRRK2-dependent consequence of IFNγ stimulation should be investigated. Additionally, although no change in GCase protein expression was detected, determination of lysosomal levels of GCase protein would be valuable given prior implication of LRRK2 in proper trafficking of GCase transporter LIMP2 to the lysosome.^66^ Overall, we find no impact of LRRK2 kinase activity on GCase activity under basal conditions. However, LRRK2 kinase activity and expression level were positively correlated with GCase activity in LWT iMGs in response to proinflammatory stimulus with IFNγ. We further assessed GCase activity in the LRRK2 variants of interest after stimulation with IFNγ. Here we observed some, although not absolute, positive correlation between increased LRRK2 kinase activity and increased GCase activity in iMGs. The p.G2019S variant did not affect kinase activity in our hands, and also did not affect GCase activity. Whereas the p.M1646T variant and correction of the p.N551K variant both led to increased kinase activity and increased GCase activity upon IFNγ stimulation. The exception to this trend was the correction of the p.R1398H variant, which led to increased kinase activity as measured by phosphorylation of Rab10, but did not increase GCase activity under inflammatory conditions, suggesting again that the p.N551K drives the protective effect seen in the p.551K-p.R1398H protective haplotype.

There is evidence that the effect of LRRK2 kinase activity on GCase activity is cell type dependent; our data implies that it is also dependent on inflammatory state, adding another layer of complexity to the interplay between these two enzymes. Inflammation, and more specifically, neuroinflammation are key hallmarks of PD. Inflammation and neuroinflammation resulting from pathogen exposure have been proposed to influence PD pathogenesis (reviewed by Tansey and colleagues),^67^ and further have been proposed to be triggered by substrate accumulation in GD and other lysosomal storage disorders.^68,69^ Here we present evidence that LRRK2 kinase activity mediates an increase in lysosomal GCase activity in response to inflammatory stimuli in microglial cells. This LRRK2-mediated increase in GCase activity under inflammatory conditions could allow microglia to better degrade excess glucosylceraminde and glucosylsphingosine, thus underlying a protective effect of LRRK2 kinase activity in *GBA1*-PD. Further investigation of the involvement of LRRK2 kinase activity in mediating GCase activity under inflammatory conditions is warranted, and a better understanding of the interaction of these two enzymes is crucial for guiding use and development of LRRK2 and GCase targeted PD therapeutics.

## Methods

### Generation and CRISPR editing of iPSC lines

The use of human iPSCs and iPSC-derived cells in this research was approved by the McGill University Research Ethics Board (IRB Study Number A03-M19-22A). PD patient derived PBMCs heterozygous for the LRRK2 p.G2019S, p.M1646T, and p.N551K-p.R1398H (protective haplotype) variants were reprogrammed as indicated in earlier studies,^48^ and provided to The Neuro’s C-BIG Open Biobank for storage and dissemination. These iPSC lines were subjected to quality control measures including karyotyping, and assessment of expression of pluripotency markers (Fig S1). LRRK2 KO and isogenic control lines with correction of LRRK2 variants were generated using CRISPR/Cas9 editing and a ddPCR based screening system. Two isogenic control lines were generated for the LRRK2 protective haplotype - one with correction of the p.N551K variant, and one with correction of the p.R1398H variant. CRISPR editing was confirmed by Sanger sequencing and LKO and isogenic control lines were subjected to the same quality control measures as their parental lines (Fig. S2). All iPSC lines used in this study are registered with the hPSCReg repository (https://hpscreg.eu/browse/provider/1355). Cell line identification numbers are as follows: LWT – IPSC0063 (https://hpscreg.eu/cell-line/CBIGi001-A), LKO – IPSC0117 (https://hpscreg.eu/cell-line/CBIGi001-A-45), p.G2019S – IPSC0006 (https://hpscreg.eu/cell-line/CBIGi006-A), p.G2019S^CORR^ – IPSC0007 (https://hpscreg.eu/cell-line/CBIGi006-A-1), p.M1646T – IPSC0017 (https://hpscreg.eu/cell-line/CBIGi013-A), p.M1646T^CORR^ – IPSC0018 (https://hpscreg.eu/cell-line/CBIGi013-A-1), p.N551K-p.R1398H – IPSC0058 (https://hpscreg.eu/cell-line/CBIGi044-A), p.N551KCORR-p.R1398H – IPSC0059 (https://hpscreg.eu/cell-line/CBIGi044-A-1), (https://hpscreg.eu/cell-line/CBIGi044-A-2). Differentiation of iMGs p.N551K-p.R1398HCORR – IPSC0060 (https://hpscreg.eu/cell-line/CBIGi044-A-2).

### Differentiation of iMGs

iPSCs were differentiated to iMGs following the protocol published by McQuade and colleagues.^49^ Briefly, iPSCs were seeded at varying densities onto matrigel (Corning 8774552) coated 6 well plates in mTESR1 (STEMCELL Technologies 85850). iPSCs were differentiated to hematopoietic precursor cells (iHPCs) using the STEMCELL Technologies STEMdiff™ Hematopoietic Kit (05310). iHPCs were collected and re-seeded in microglial differentiation media (MDM) + three cytokine cocktail (M-CSF, IL-34, and TGF-β), onto matrigel coated 6 well plates on days 10 and 12 of iHPC differentiation. Cells were supplemented with MDM + three cytokine cocktail every other day to mediate microglial differentiation. A full media change was performed on day 12, and a media change to MDM + five cytokine cocktail (M-CSF, IL-34, TGF-β, CD200, and CX3CL1) on day 25. Cells were supplemented with MDM + five cytokine cocktail on day 27 and considered mature on day 28. Human recombinant M-CSF (300-25), IL-34 (200-34), TGF-β (100-21), and CX3CL1 (300-31) purchased from Peprotech. Human recombinant CD200 (BP004) purchased from Bonopus Bioscience. For immunofluorescence, PFB-FDGlu, and DQ-BSA assays mature iMGs were replated. iMGs were dissociated by scraping in PBS, and replated onto 12 mm diameter glass coverslips at a density of 100,000 cells per coverslip, or 96-well Flat Clear Bottom Black Polystyrene TC-treated Microplates (Corning 3904) without Matrigel coating at a density of 20,000 cells per well. iMGs were supplemented with MDM + five cytokine cocktail every other day and assayed five days after replating.

### Flow cytometry detection of CD45 and CD11b

Mature iMGs were collected; non-adherent cells are collected with media, while adherent cells are scraped in PBS. iPSCs were harvested in a single-cell suspension using accutase (STEMCELL Technologies 07922) dissociation. iPSCs and iMGs were stained with Live/Dead fixable aqua stain (ThermoFisher L34965) in PBS for 30 minutes, washed with FACS buffer (1X PBS, 1% FBS, 0.1% NaN_3_) and blocked with TrueStain blocking reagent (Biolegend 422302) for 10 minutes. Cells were stained with anti-CD45 – AlexaFluor 700 (1/40 Biolegend 304024) and CD11b – PE (1/20 BD Biosciences 555388) fluorophore-conjugated antibodies for 15 minutes, washed and resuspended in FACS buffer for analysis. iPSCs and iMGs were analyzed on Thermo Attune NxT cytometer (Thermo) equipped with 405, 488, and 561 nm lasers and 610/20, 620/15, 530/30, and 525/50 filters (NeuroEDDU Flow Cytometry Facility, McGill University). For each sample, 50,000 events were collected and single, live cells were subsequently gated for CD45 and CD11b. Data were analyzed using FlowJo (BD Biosciences).

### Immunofluorescence staining of Iba1 and PU.1

iMGs replated onto coverslips were fixed with 4% paraformaldehyde for 20 minutes. iMGs were washed three times with PBS, permeabilized with 0.2% triton-X 100 in PBS for 10 minutes, and blocked with 5% normal donkey serum (NDS) in PBS for one hour at room temperature. iMGs were incubated with primary antibodies against Iba1 (1/1,000, Synaptic Systems 234 009) and PU.1 (1/500, Cell Signalling 2266S) diluted in 5% NDS in PBS overnight at 4_°_C. iMGs were then washed with PBS three times, and incubated with a 1/500 dilution of donkey anti rabbit AlexaFluor 647 conjugated secondary antibody (Invitrogen A-32795), a 1/500 dilution of donkey anti chicken AlexaFluor 488 conjugated secondary antibody (Invitrogen A-78948) and 1 mg/mL Hoechst in 5% NDS in PBS for one hour at room temperature. iMGs were washed three times in PBS, mounted onto slides using Aqua-Poly/Mount (Polysciences Inc. 18606-20) and imaged on a Leica SP8 confocal microscope.

### iMG treatments

For LRRK2 inhibition, iMGs were treated with 100 nM MLi-2 (Tocris 5756) or 500 nM PF-475 (MedChemExpress HY-12477) for six hours. For IFNγ stimulation experiments, iMGs were pretreated with DMSO or 100 nM MLi-2 for one hour. iMGs were then treated with 20 ng/mL IFNγ (Peprotech 300-02) or vehicle and DMSO or 100 nM MLi-2 for 24 hours. GCase inhibition was achieved with 25 nM Conduritol B Epoxide (Sigma C5424) treatment for 25 hours.

### Western blotting

Mature iMGs were collected, and resuspended in lysis buffer (50 mM Tris-HCl pH 7.4, 1% v/v Triton-X 100, 10% Glycerol, 1X Halt phosphatase inhibitor cocktail (Thermo 78428), 0.1 mg/mL microcystin-LR (Enzo Life Sciences ALX-350-012), 1X Complete protease inhibitor cocktail (Sigma 11873580001)). Samples were then incubated at 4°C with rotation for 30 minutes, followed by three rounds of sonication in a water bath (30 seconds in water, 30 seconds on ice). Finally, samples were centrifuged at 20 800 x g for 20 minutes at 4°C. Supernatants were collected and protein concentration measured using Detergent Compatible Protein Assay (Bio-Rad 5000112). Samples were prepared with a total of 25 μg of protein, and resolved on 4-15% or 15% SDS-PAGE gels, then transferred to PVDF using a Trans-Blot Turbo System (Bio-Rad). Membranes used for detection of α-synuclein were fixed in 4% paraformaldehyde, 0.1% glutaraldehyde for 30 minutes at room temperature, and washed three times in TBS with 0.1% Tween-20 (TBS-T). Membranes were blocked in 5% bovine serum albumin (BSA) in TBS-T. Membranes were incubated with primary antibodies diluted in 5% BSA in TBS-T overnight at 4°C with shaking. Primary antibodies used and dilutions are as follows: LRRK2 pS1292 1/200 (Abcam ab203181), LRRK2 pS935 1/500 (Abcam ab133450), total LRRK2 1/500 (Abcam ab133474), Rab10 pT73 1/500 (Abcam ab230261), total Rab10 1/500 (Cell Signaling 8127S), Rab12 pS106 1/1,000 (Abcam ab256765), total Rab12 1/500 (Proteintech 18843-1-AP), GCase 1/1,000 (Abcam ab55080), α-synuclein 1/1,000 (BD Biosciences 610787), LAMP1 1/1,000 (Cell Signaling 9091S), α-actin 1/50,000 (Millipore MAB1501). Membranes were washed three times with TBS-T, incubated with HRP-conjugated secondary antibodies diluted at 1/5,000 in 5% BSA in TBS-T for one hour at room temperature, and washed as before. Western blots were visualised with Clarity Western ECL Substrate or Clarity Max Western ECL Substrate (Bio-Rad 170–5061, 170-5062) on a ChemiDoc MP Imaging System (Bio-Rad). Analysis performed using FIJI with α-actin as a loading control.

### PFB-FDGlu GCase assay

Replated iMGs were treated with LRRK2 inhibitor and/or IFNγ as described above. iMGs were stained with lysotracker deep-red (1:20,000, Invitrogen L12492) for 30 minutes. Media was then exchanged for fluorobrite imaging media (Thermo A1896701) with 37.5 uM PFB-FDGlu (Invitrogen P11947). LRRK2 inhibition was continued throughout. iMGs were imaged on an Opera Phenix high-content confocal microscope (Revvity) every 10 minutes for a total of 140 minutes. Columbus software was used to identify cells (using lysotracker signal) for quantification of mean PFB-FDGlu fluorescence per cell, and mean total lysotracker area per cell.

### DQ Red BSA Lysosomal Proteolysis Assay

Replated iMGs were treated with LRRK2 inhibitor and/or IFNγ as described above. Concomitant with IFNγ treatment cells were loaded with 1 μg/mL DQ Red BSA (Invitrogen D12051) for 24 hrs. iMGs were stained with lysotracker deep-red (1:20,000, Invitrogen L12492) for 30 minutes. Media was exchanged for fluorobrite imaging media and cells were imaged on an Opera Phenix high-content confocal microscope. Columbus software was used to identify cells (using lysotracker signal) and mean DQ-BSA fluorescence per cell was quantified and normalized to mean total lysotracker area per cell.

### pHLys Green Lysosomal pH Assay

Lysosomal pH was assayed using the Lysosomal Acidic pH Detection Kit-Green/Deep Red (Dojindo L286-10). Replated iMGs were treated with LRRK2 inhibitor, MLi-2 and/or IFNγ as described above. iMGs were then loaded with LysoPrime Deep Red (1:1,000) for 30 minutes. iMGs were then washed once with Hank’s balanced salt solution (HBSS, Gibco 14175095), and loaded with pHLys Green diluted 1:1,000 in HBSS for 30 minutes. pHLys Green was removed and cells were imaged in HBSS on an Opera Phenix high-content confocal microscope. Columbus software was used to identify cells (using LysoPrime Deep Red signal) and mean pHLys Green fluorescence per cell was quantified and normalized to mean total LysoPrime Deep Red area per cell.

### Statistical Analysis

Statistical analysis was conducted in GraphPad Prism9 software. Biological replicates are defined as experiments conducted using distinct differentiations from iPSCs to iMGs. All plots depict the mean value across biological replicates -/+ standard deviation. Statistical comparisons were computed using One way ANOVA with Bonferroni post test for western blotting experiments, DQ-BSA, and pHLys assays. Repeated Measures One way ANOVA with Tukey post test was utilized for PFB-FDGlu assays. Significance levels are depicted in figure legends.

## Author Contributions

EJM generated p.M1646T and protective haplotype isogenic correction cell lines, differentiated iMGs, designed and performed experiments, analysed data, and prepared the manuscript. CXQC and NA performed quality control experiments on iPSC lines used in this study. ED designed CRISPR editing strategies and ddPCR screening strategies to generate isogenic correction lines, and generated the p.G2019S isogenic correction line. ZY designed CRISPR KO strategy and generated the LKO line. TM, KS, and ZG-O designed experiments. EAF designed experiments, supervised the project, and prepared the manuscript.

## Acknowledgements

Thanks to Wolfgang Reintsch, and Julien Sirois for training and support with high content imaging and flow cytometry experiments, respectively. Thanks to Marie-France Dorion for training and advice on differentiation and characterization of iPSC-derived microglia. Thanks to Jace Jones-Tabah, Roxanne Larivière, and Andrea Krahn for establishing and optimizing live-cell GCase, lysosomal proteolytic and pH assays. EJM has been supported by a Fonds de Recherche du Québec-Santé Doctoral Fellowship and Jeanne Timmins Costello Fellowship awarded by the Integrated program in Neuroscience at McGill University. This work was funded by grants from The Michael J. Fox Foundation for Parkinson’s Research (MJFF-019045) and from the Canadian Institutes of Health Research (FDN-154301). G-Can (GBA1 Canada) Initiative, an open-science collaborative initiative aimed at addressing GBA1 associated neurodegeneration, has contributed to this research. G-Can is supported by The Hilary & Galen Weston Foundation, J. Sebastian van Berkom and Ghislaine Saucier and Silverstein Foundation. EAF is supported by a Canada Research Chair (Tier 1) in Parkinson’s disease.

## Data Availability

All data generated or analysed during this study are included in this published article.

## Figure Captions

**Figure S1.**
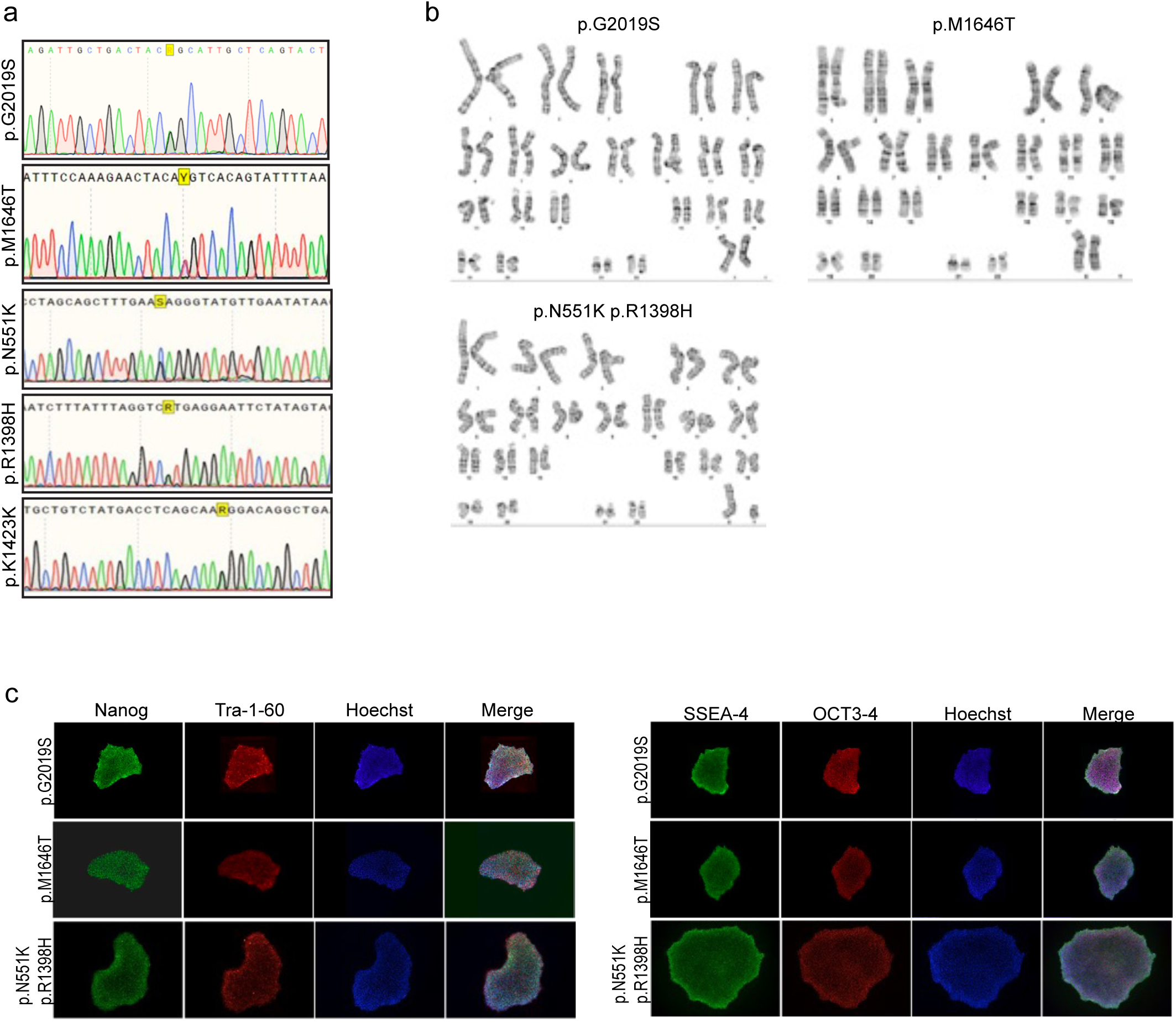
Sequencing and quality control of PD patient-derived LRRK2 variant iPSC lines used in this study. **a** Sanger sequencing confirmation of heterozygous LRRK2 variants. **b** Karyotype analysis shows no chromosomal abnormalities in iPSC lines. **c** Expression of pluripotency markers by IF staining.

**Figure S2.**
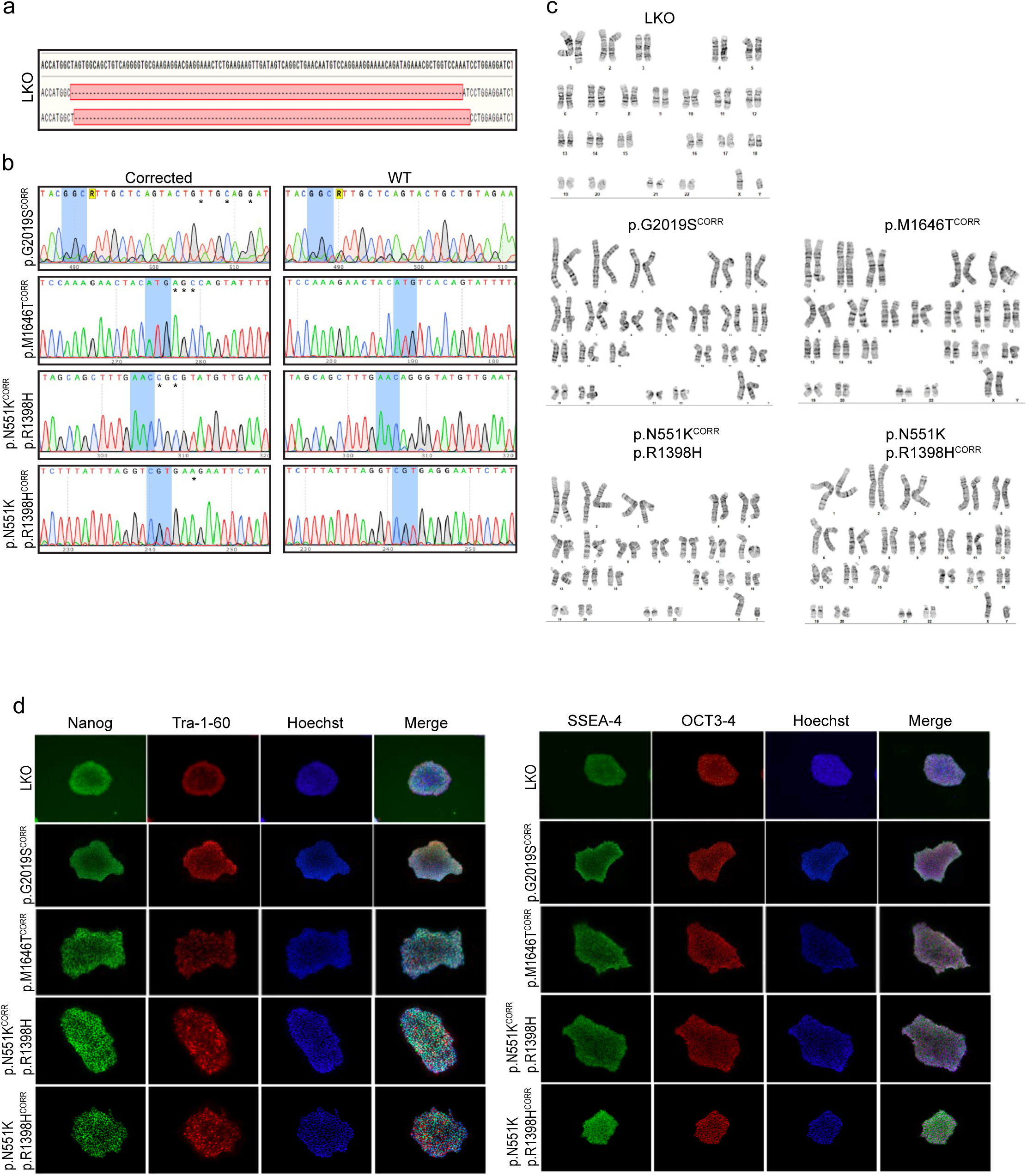
Sequencing and quality control of CRISPR-edited LKO and isogenic control iPSC lines used in this study. **a** Sanger sequencing confirms disruption of the LRRK2 gene by a 106 base-pair or 107 base-pair deletion, both resulting in frame shift mutations. **b** Sanger sequencing confirmation of correction of heterozygous LRRK2 variants, and introduction of PAM disrupting silent mutations. Corrected sequence illustrates the CRISPR-edited variant allele now corrected, with additional silent mutations indicated by *. WT sequence is that of the non-edited, non-variant allele. **c** Karyotype analysis shows no chromosomal abnormalities in iPSC lines. **d** Expression of pluripotency markers by IF staining.

**Figure S3.**
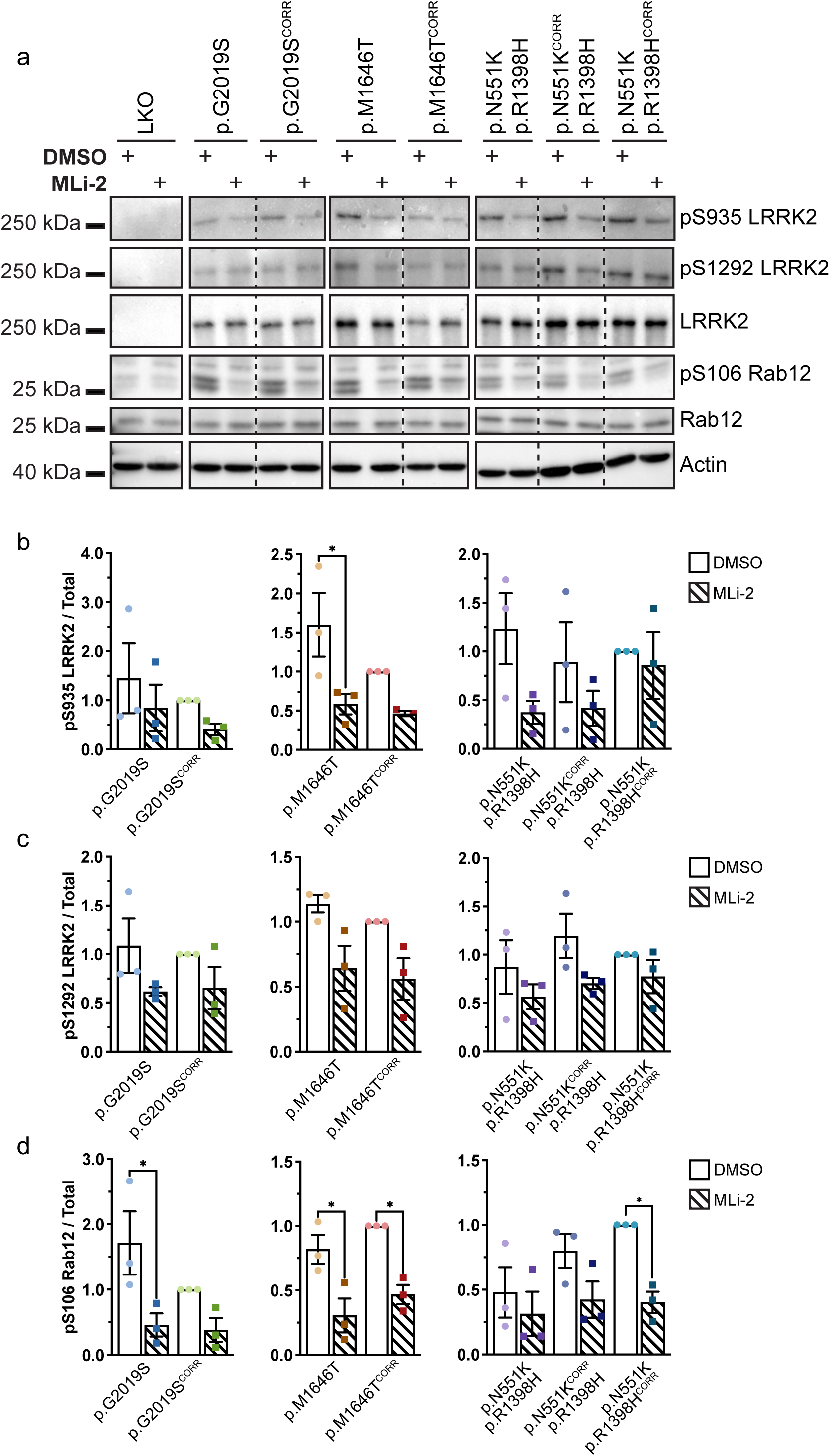
Rab12 and LRRK2 phosphorylation unchanged in LRRK2 variant iMGs. **a** LRRK2 and Rab12 phosphorylation in LRRK2 variant iMGs with or without 6 hr 100 nM MLi-2 treatment as measured by WB. n = 3 **b-d** Quantification of WB of phosphorylated LRRK2 or Rab12 normalized to total LRRK2 or Rab12, levels normalized to isogenic control. **b** Quantification of pS1292 LRRK2. **c** Quantification of pS935 LRRK2. **d** Quantification of pS106 Rab12. One Way ANOVA with Bonferroni post-hoc test * p < 0.05, ** p < 0.01, **** p < 0.0001

**Figure S4.**
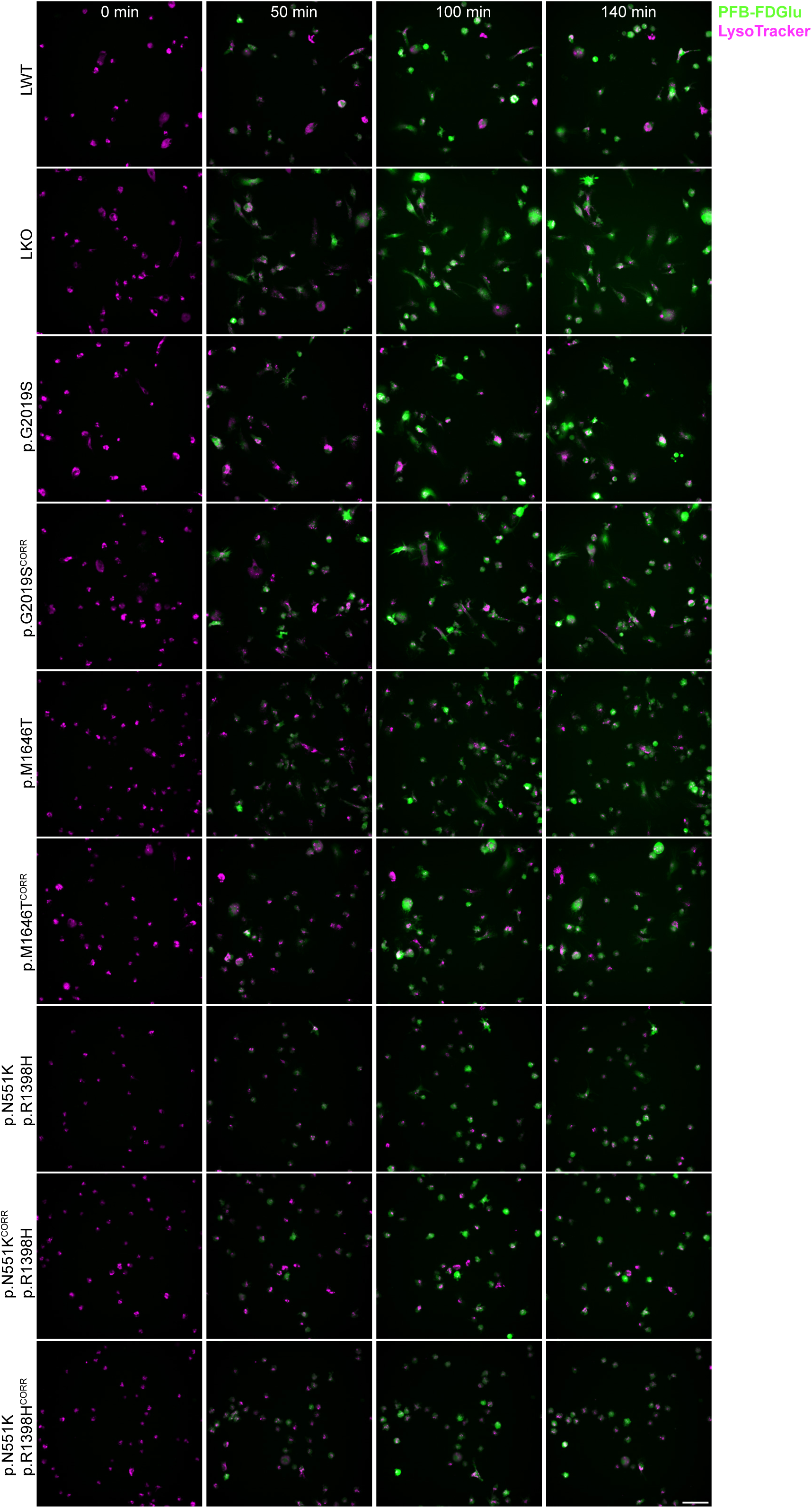
PFB-FDGlu GCase assay images from LRRK2 variant iMGs stained with lysotracker deep-red 0, 50, 100, and 140 minutes after dye-loading. Scale bar 50 μm.

**Figure S5.**
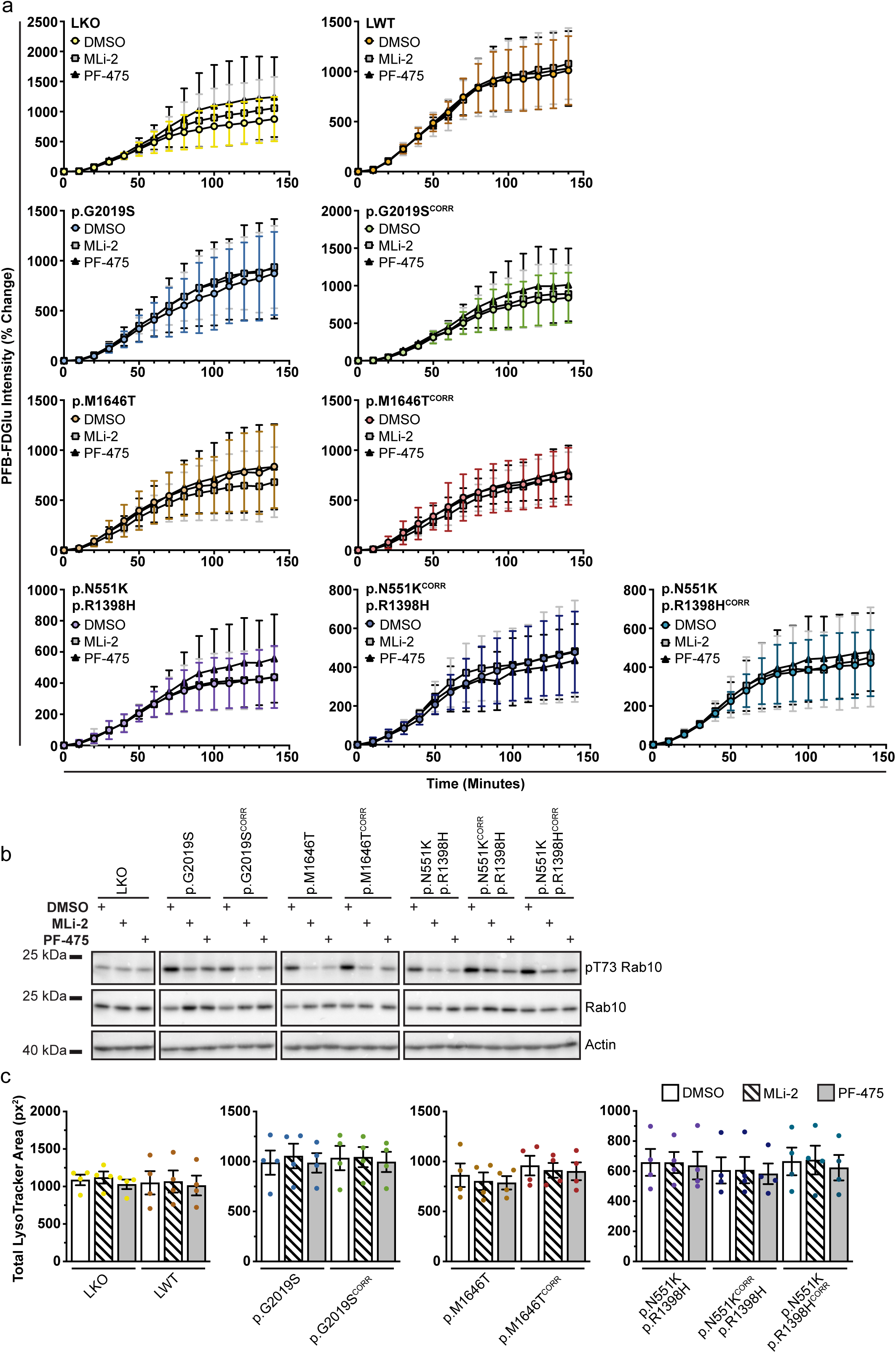
LRRK2 inhibition leads to Rab10 dephosphorylation but has no effect on GCase activity. **a** Percent change in PFB-FDGlu fluorescence. n = 4. **b** Rab10 phosphorylation is decreased to a similar extent by 6 hr 100 nM MLi-2 or 500 nM PF-475 treatment. n = 2 **c** Mean total lysotracker fluorescence area per cell (px^2^) at baseline (0 minutes). **a & c** Repeated Measures One Way ANOVA Tukey post-hoc test * p < 0.05, ** p < 0.01, **** p < 0.0001

**Figure S6.**
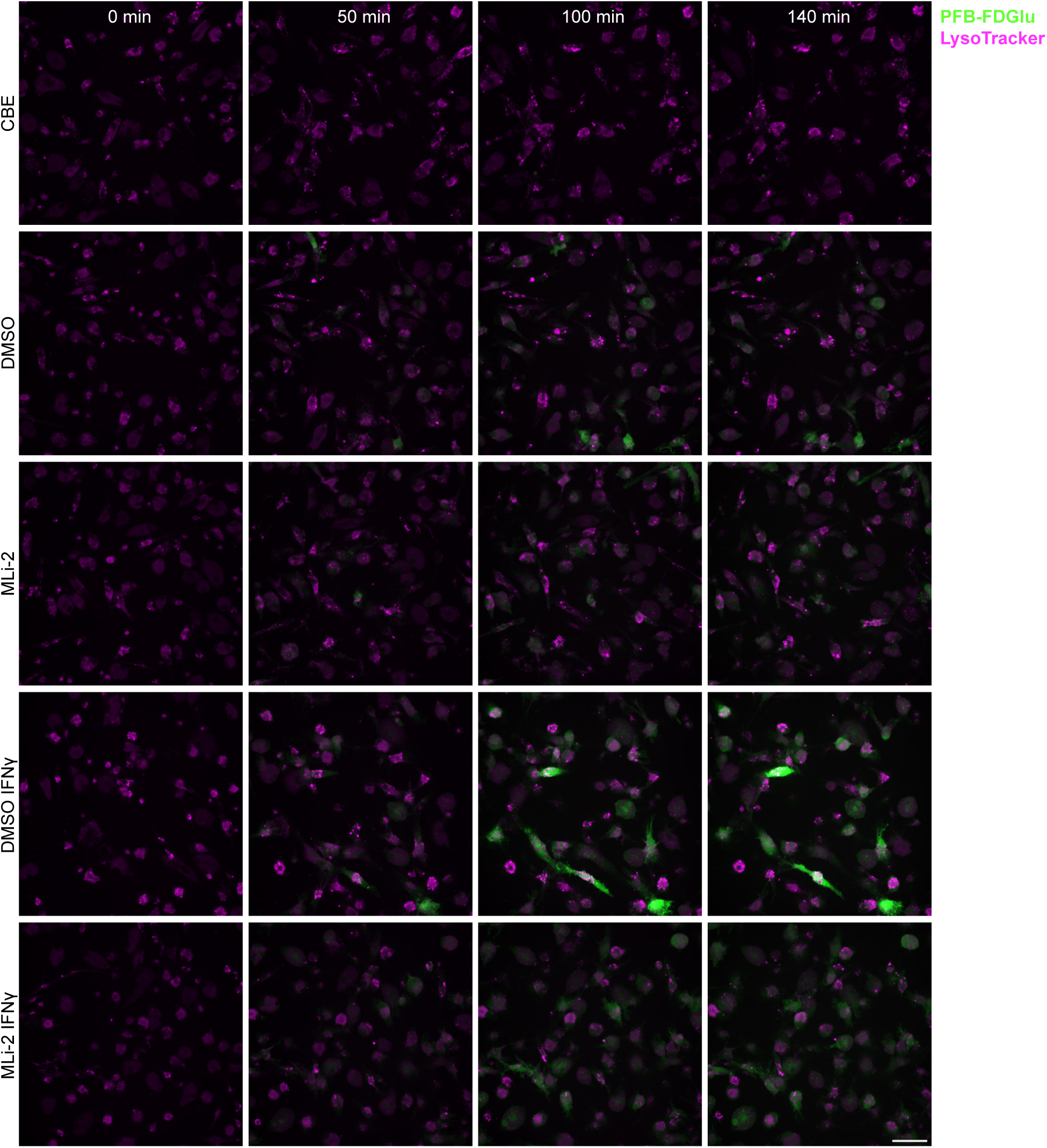
PFB-FDGlu GCase assay images from LWT iMGs treated with 20 ng/mL IFNγ and 100 nM MLi-2 stained with lysotracker deep-red 0, 50, 100, and 140 minutes after dye-loading. Scale bar 50 μm.

**Figure S7.**
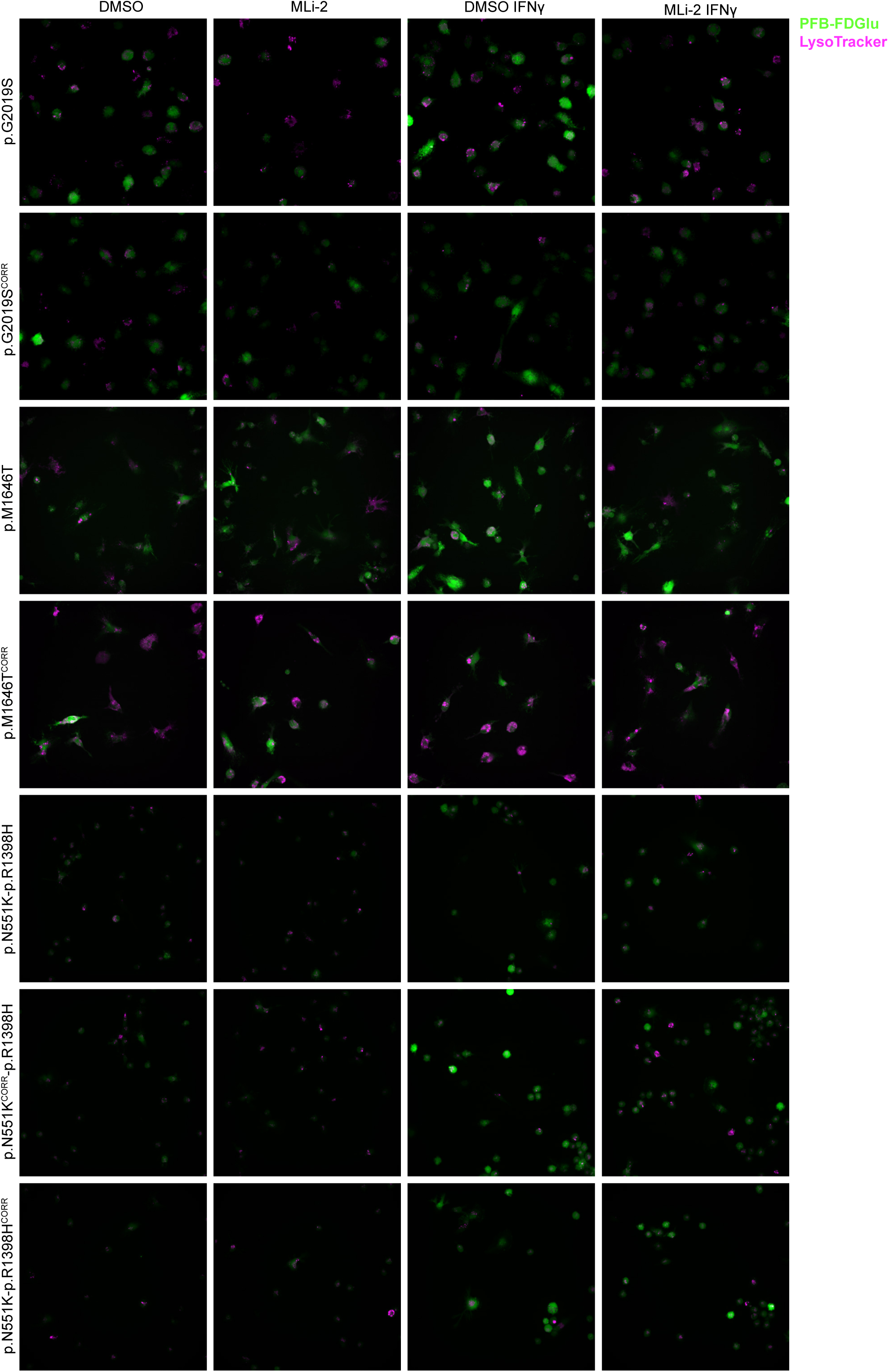
PFB-FDGlu GCase assay images from LRRK2 variant and isogenic control iMGs with or without 20 ng/mL IFNγ and/or 100 nM MLi-2, stained with lysotracker deep-red 140 minutes after dye-loading. Scale bar 50 μm.

